# Broad Spectrum Proteomics Analysis of the Inferior Colliculus following Acute Hydrogen Sulfide Exposure

**DOI:** 10.1101/237370

**Authors:** Dong-Suk Kim, Poojya Anantharam, Andrea Hoffmann, Mitchell L. Meade, Nadja Grobe, Jeffery M. Gearhart, Elizabeth M. Whitley, Belinda Mahama, Wilson K. Rumbeiha

**Author notes:** Corresponding Author’s Contact Information. Present addresses: Poojya Anantharam: Medical Countermeasures, MRI Global, Kansas City, MO, US. Belinda Mahama: Neuroscience, Brown University, Rhode Island, Providence, US.

## Abstract

Acute exposure to high concentrations of H_2_S causes severe brain injury and long-term neurological disorders. The mechanisms of H_2_S-induced neurodegeneration are not known. To better understand the cellular and molecular mechanisms of H2S-induced neurodegeneration we used a broad-spectrum proteomic analysis approach to search for key molecules in H2S-induced neurotoxicity. Mice were subjected to acute whole body exposure of up to750 ppm of H_2_S. The H2S-treated group showed behavioral motor deficits and developed severe lesions in the inferior colliculus (IC), part of the brainstem. The IC was microdissected for proteomic analysis. Tandem mass tags (TMT) liquid chromatography mass spectrometry (LC-MS/MS)-based quantitative proteomics was applied for protein identification and quantitation. LC-MS/MS was able to identify 598, 562, and 546 altered proteomic changes for day 1 (2 h post H_2_S exposure), day 2, and day 4 of H_2_S exposure, respectively. Mass spectrometry data were analyzed by Perseus 1.5.5.3 statistical analysis, and gene ontology heat map clustering. Quantitative real-time PCR was used to confirm some of the H_2_S-dependent proteomics changes. Taken together, acute exposure to H_2_S induced behavioral motor deficits along with progressive neurodegeneration including disruption of several biological processes in the IC such as cellular morphology, energy metabolism, and calcium signaling. The obtained broad-spectrum proteomics data may provide important clues to elucidate mechanisms of H_2_S-induced neurotoxicity.

**Highlights:**

- Mice exposed to H_2_S recapitulated H_2_S-induced neurotoxicity manifested in humans.
- The IC in the mouse brain is the most sensitive to H_2_S-induced neurodegeneration.
- Proteomic expressions of key proteins were validated at transcription level.
- Several biological pathways were dysregulated by H_2_S exposure.

## 1. Introduction

Hydrogen sulfide (H_2_S) is a highly neurotoxic colorless gas with a “rotten egg” odor (Chou *et al.*, 2016). It is as an environmental pollutant that causes occupational hazards in a variety of industrial processes including the oil and gas industry, intensive animal farming operations, sewer and waste water treatment plants, pulp and paper plants, and gas storage facilities, among several others (Chou *et al.*, 2016). It has been estimated that there are more than 1,000 reports of human exposures to H_2_S each year in the United States (Chou *et al.*, 2016). Besides accidental H_2_S poisoning in industrial settings, intentional acute exposure to high concentrations of H_2_S as a means of suicide has been increasingly observed in Western and Asian societies (Morii *et al.*, 2010; Reedy *et al.*, 2011). The raw chemical ingredients used to generate H_2_S in such circumstances are readily accessible in local stores (Morii *et al.*, 2010). More significantly, H_2_S is listed as a high priority chemical by the US Department of Homeland Security because of its potential to be misused in nefarious acts such as chemical terrorism, particularly in confined spaces such as the underground transit systems. Although H_2_S is toxic at high concentrations, it is also beneficial at physiologic concentrations. For this reason, there is also tremendous interest in potential therapeutic applications of H_2_S for treatment of a number of human disease conditions. Proposed pharmacological uses of H_2_S include ulcer treatment, ischemic reperfusion injury, as an anti-inflammatory for treatment of Crohn’s disease, endotoxin induced inflammation, arterial hypertrophy, visceral pain, Parkinson’s disease and cancer among others (Szabo, 2007; Popov, 2013). Furthermore, H_2_S was shown to activate nitric oxide (NO) synthesis through induction of endothelial nitric oxide synthase. Similar to NO, H_2_S serves as a gasotransmitter and signaling molecule in the CNS and regulating vasodilation (Tan *et al.*, 2010).

Like most toxicants, the toxicity of H_2_S exposure is dose-dependent, but H_2_S characteristically has a steep-dose response curve (Guidotti, 1996). At concentrations below 30 parts per million (ppm), H_2_S causes headache, coughing and throat irritation. At 150 ppm H_2_S induces olfactory fatigue and temporary loss of smell after 5-60 min of exposure (Wasch *et al.*, 1989). Concentrations higher than 500 ppm cause headache, dizziness, unconsciousness, and respiratory failure. At high concentration above 1,000 ppm H_2_S can lead to immediate loss of consciousness, commonly called “knock-down”, and ultimately seizures, and sudden death (Chou *et al.*, 2016). Typically, acute exposure to high concentrations of H_2_S is associated with high mortality withing a few hours post-exposure (McCabe and Clayton, 1952). Timely rescue of victims can prevent disaster and allow victims to fully recover. Currently, there is no effective antidote to treat victims of H_2_S gas exposure and the mortality rate remains high (Lindenmann et al.; Herbert, 1989; Vicas, 1989; Smith, 1997; Lindenmann *et al.*, 2010). In addition, delayed neurological disorders, which can last for many years, are commonly reported in survivors of acute H_2_S exposures (Matsuo *et al.*, 1979; Parra *et al.*, 1991; Tvedt *et al.*, 1991a; Tvedt *et al.*, 1991b; Snyder *et al.*, 1995; Kilburn, 2003; Woodall *et al.*, 2005). These delayed neurological sequelae are incapacitating, and require prolonged medical attention that lacks defined medical interventions.

The development of effective therapeutics requires a good understanding of the molecular mechanisms and pathways of H2S-induced neurodegeneration and neurological sequelae. These mechanisms remain largely unknown. There is an acute need for countermeasures for treatment of mass civilian casualties of acute H_2_S poisoning in the field, such as following catastrophic industrial meltdowns or intentional terrorist activities. Elucidating molecular mechanisms underlying H_2_S-induced neurotoxicity is essential in developing suitable targeted therapeutics to counter both acute and delayed neurotoxic effects of H_2_S poisoning in humans.

We recently developed a relevant inhalational animal model that exhibits the clinical, pathological, and motor behavioral symptoms of H_2_S exposed survivors (Anantharam *et al.*, 2017). The objective of this study was to build on our previous work and investigate proteomic changes and altered gene expression in a mouse model of acute H_2_S-exposure to identify novel toxic mechanisms. In prior studies, we discovered that the central inferior colliculus (IC) region of the brainstem is the most sensitive brain region to H_2_S-induced neurodegeneration (Anantharam *et al.*, 2017). Consequently, we focused on the IC in this proteomic study, although we have also observed histopathological changes in other parts of the brain such as the thalamus and cortex (Anantharam *et al.*, 2017). This study demonstrated, for the first time, that H_2_S exposure induces significant proteomic changes in the IC, which may play an important role in execution of H_2_S-induced neurotoxicity.

## 2. Materials and Methods

### 2.1 Chemicals

Hydrogen sulfide gas was purchased from Airgas (Radnor, PA). RNeasy mini kit was purchased from Qiagen (Germantown, MD). High Capacity cDNA RT kit were purchased from ThermoFisher Scientific (Waltham, MA). RT^2^ SYBR Green ROX qPCR Mastermix and primers for Gapdh were purchased from Qiagen (Valencia, CA).

### 2.2 Animals and treatment

This study was approved by Iowa State University Animal Care and Use Committee. Seven-to eight-week-old male C57 BL/6J mice were housed at room temperature of 20 – 22 °C under a 12-h light cycle, and a relative humidity of 35 - 50 %. Protein Rodent maintenance diet (Teklad HSD Inc, WI, US) and water were provided *ad libitum*. Prior to exposure on day 1, all mice were acclimated to breathing air for 40 min on each of two days preceding day 1 when mice were first exposed to H_2_S. Mice were exposed by whole body inhalation exposure either to normal breathing air from a tank under pressure (Control) or to 650 - 750ppm H_2_S (H_2_S-treated) once or for short-term daily exposures lasting for up to 6 days. The exposure paradigm is summarized in Fig 1. On day 1, mice were exposed to H_2_S of 650 - 750 ppm for 40 min only. Those mice terminated on day 1 were euthanized 2 h post H_2_S-eposure. The remaining groups of H_2_S-exposed mice were exposed daily to 765 ppm H_2_S for 15 min. only. In this regard, mice euthanized on day 1 received only 1 acute exposure, those euthanized on day 2 received 2 acute exposures of H_2_S and those euthanized on day 4 received 4 acute exposures to H_2_S. Negative controls were exposed to normal breathing air daily and euthanized on day 4. Mice were euthanized by decapitation 2 h after the last exposure to H_2_S on days 1, 2 and 4 for proteomics analysis and quantitative gene expression analysis by real-time PCR and on days 1, 3 and 7 for histopathology evaluation. Following decapitation, brains were immediately removed from the skull. The IC were microdissected on ice and immediately flash frozen using liquid nitrogen, and stored at −80°C until further use. Animals were cared for in accordance with the Institutional Animal Care and Use committee guidelines.

**Figure 1.**
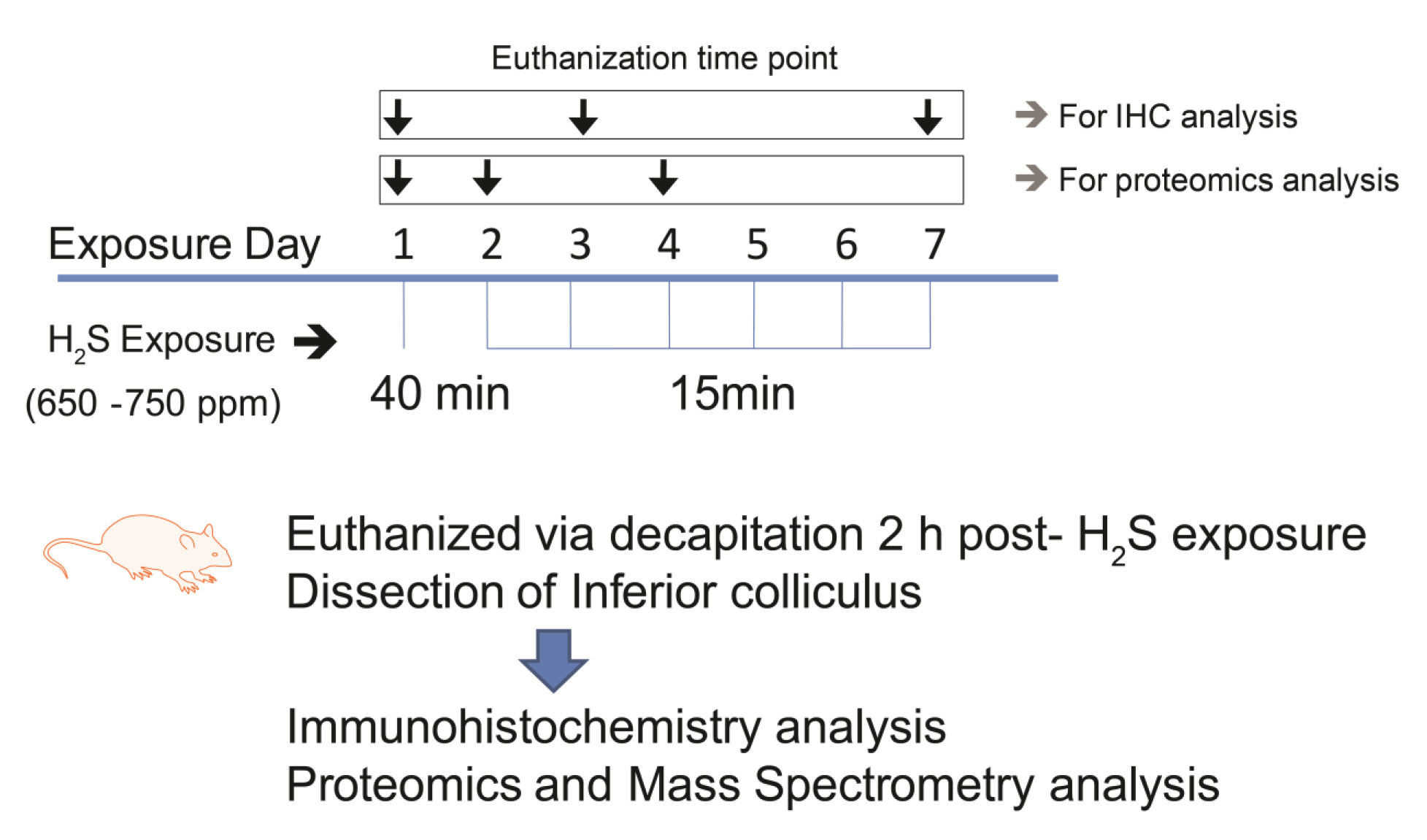
Acute exposure paradigm of hydrogen sulfide on C57 black mice. Mice were exposed to 765 ppm of hydrogen sulfide in a chamber for 40 min either once only (day 1) and for 15 min on the following days up to day 7. Mice were sacrificed 2 h post-H_2_S exposure on specified days of the study. Negative control mice were exposed to breathing air from a cylinder daily up to day 7. Separate groups of mice were sacrificed on days 1, 3, and 7 for immunohistochemistry. Groups of mice for proteomics studies were sacrificed on days 1, 2 and 4. (One column figure)

### 2.3 Behavioral assessment

The VersaMax open field test was used to assess motor deficits induced by H_2_S. We used this test because previous studies in the lab had indicated it to be sensitive to acute H_2_S intoxication in this mouse model (Anantharam *et al.*, 2017). Spontaneous activity was measured using an automated computer device (Model RXYZCM-16; Accuscan, Columbus, OH, USA). The activity chamber’s dimensions are 40 × 40 × 30.5 cm, and it is made of Plexiglas with a Plexiglas lid. The lid has holes for ventilation. Data was analyzed using VersaMax Analyzer (Model CDA-8; Accuscan). Mice were placed in the activity chamber 2 min prior to recording for 10 min, for acclimation. Vertical activity, horizontal activity, and distance travelled were measured.

### 2.4 Histopathology

Mice were euthanized 24 h after the last H_2_S exposure. They were deeply anesthetized with a cocktail of 100 mg/kg bw ketamine and 10 mg/kg bw xylazine. The thoracic cavity was opened to expose the heart and fresh 4 % paraformaldehyde (PFA, pH 7.4) was injected through the left ventricle. After perfusion, the calvarium was opened and brains were post-fixed in 4 % PFA for 24-48 h before removal from the skull. Brains were processed in paraffin, sectioned at 5 microns, and stained with hematoxylin and eosin, and examined microscopically. Neurons were stained with NeuN antibody (ab177487, Abcam, Cambridge, MA) using an indirect immunostaining protocol. Diaminobenzidine was used as chromogen. Stained sections were examined using a Nikon Eclipse Ci-L microscope equipped with a DS-Fi2 camera. For image analysis for the quantification of neurons, NeuN-positive cells (sites of DAB chromogen deposition) were enumerated in each of five 400X photomicrographs of the IC from mice exposed to H_2_S or breathing air, and the mean number of NeuN-positive cells/mouse were compared between groups.

### 2.5 Mass Spectrometry Analysis

Samples were processed according to a previously published method with some modifications (Meade *et al.*, 2015). Briefly, IC tissues were placed in 100μl of urea lysis buffer and homogenized with a handheld pestle homogenizer. Protein samples were further reduced and alkylated. Small aliquots of each sample were taken to measure protein concentration using the Bradford assay from Bio-Rad (Hercules, CA). The remaining samples were diluted and trypsin-digested overnight, followed by desalting using a C18 peptide trap from Michrom (Auburn, CA). The desalted samples were vacufuged prior to individual sample labeling using TMT-6plex labels according to the manufacturers’ instructions (Thermo FisherScientific, Waltham, MA). Labeled samples of each exposure group were combined with the control for sample comparison (Table 1). Peptides were separated on a Waters BEH C18 capillary column prior to online analysis using a 240-min linear increasing gradient of acetonitrile with 0.1% formic acid. Following elution from the column, ions were generated using 2.6 kV on a taper tip in a New Objective nanosource and entered into an LTQ-OrbitrapVelos mass spectrometer (Thermo Fisher, San Jose, CA). A full scan was taken in the LTQ, followed by data-dependent MS/MS analysis of the top 6peaks. MS/MS analysis included collision-induced dissociation (CID) in the LTQ for structural information and higher-energy collisional dissociation (HCD) in the Orbitrap for quantitation. MS/MS data were aligned and quantitated using MaxQuant1.5.4.1 (Cox and Mann, 2008) Analytics Platform with PTXQC (Bielow *et al.*, 2016) quality control data management. Peptide alignment was executed with the mouse Uniprot protein database UP000000589_10090.fasta; enzyme: trypsin; carboxymethyl (C), and oxidation (M), FDR <1% based on peptide q-value under standard settings (see supplemental data). The secondary computational analysis was executed using Perseus 1.5.5.3 (Tyanova *et al.*, 2016) for statistical rendering, and web-based software Morpheus for heatmap rendering. Prior to analysis of experimental samples, small aliquots of individually labeled control samples were analyzed to determine individual variations in controls. Due to low variability in the controls, a pooled control setting was used. Sample analysis included modification of the MaxQuant protein output list by normalization of individual TMT reporter ion intensities by division through the median intensity followed by determination of fold TMT expression ratios by division of individual experimental TMT values (126,127,128,129,and 130) through the TMT value (131) of the pooled control.

**Table 1.** Animal exposure paradigm for mass spectrometry and distribution of TMT sample labeling. GA= group A; GB= group B; GC=group C; GD= group D. (two column table)

| Group | Total Mice/<br>Animal ID | TMT<br>Label | Treatment | Exposure/<br>Dose | Euthanasia |
| --- | --- | --- | --- | --- | --- |
| GA<br>Control | (n=5)<br>1,2,3,4,5,<br>pooled | 131 | 40 min day 1<br>15 min day 2<br>15 min day 3<br>15 min day 4<br>+ 2 h recovery | Inhalation<br>/<br>air | Day 4 |
| GB<br>1<br>exposure | (n=5)<br>1<br>2<br>3<br>4<br>5 | 126<br>127<br>128<br>129<br>130 | 40 min day 1<br>+ 2 h recovery | Inhalation<br>/<br>650 - 700<br>ppm H<sub>2</sub>S | Day 1 |
| GC<br>2<br>exposures | (n=5)<br>1<br>2<br>3<br>4<br>5 | 126<br>127<br>128<br>129<br>130 | 40 min day 1<br>15 min day 2<br>+ 2 h recovery | Inhalation<br>/<br>650 – 700<br>ppm H<sub>2</sub>S | Day 2 |
| GD<br>4 | (n=4)<br>1 | 126 | 40 min day 1<br>15 min day 2 | Inhalation<br>/ | Day 4 |
| exposures | 2 | 127 | 15 min day 3 | 650 – 700 |  |
|  | 3 | 128 | 15 min day 4 | ppm H<sub>2</sub>S |  |
|  | 5 | 130 | + 2 h recovery |  |  |

### 2.6 Perseus Statistical Analysis

The previously modified Maxquant protein list was entered into the Perseus 1.5.5.3 software program followed by annotating Maxquant defined Swissprot protein accession numbers with mouse gene identifiers. Statistical analysis included one-sample t-tests of fold expression values by considering the deviation of samples from 1 fold expression (no change in TMT reporter ion intensities vs. control intensity values) using p<0.05 as a criterion for significance. Scatter Plots were established by plotting one-sample t-test difference fold protein expression vs. −log one-sample t-test p value fold protein expression according to the associated Perseus tool set.

### 2.7 Morpheus Heatmap Rendering

The Perseus processed protein list containing mouse gene identifiers, protein names and gene ontology (GO) biological process, one-sample t-test, levels of significantly and non-significantly modulated proteins was entered into the web-based Morpheus heatmap-rendering tool (https://software.broadinstitute.org/morpheus/). A gradient coloring scheme was applied to display upregulated (above 1.2 fold expression (in log2 display above 0.263), red color) vs. downregulated (below 0.83 fold expression (in log2 display below −0.263) blue color) protein nodes. White color nodes represent changes of non-modulated nodes to control levels (between 0.83 and 1.2 fold expression). The heatmap was hierarchically clustered by Euclidean distance using row average linkage and grouping of rows by GO biological process.

### 2.8 Validation of gene expression via quantitative real-time RT-PCR

After mice were exposed to H_2_S, tissues were dissected and immediately stored at −80 °C till analysis. Total RNA was extracted from frozen tissues using the RNeasy^®^ Plus Mini kit with treatment of DNase I according to the manufacturer’s protocol. Validated primers for Gapdh (Qiagen, #PPR57734E) were used as the housekeeping gene controls. The threshold cycle (C_t_) was calculated from the instrument software, and fold change in gene expression was calculated using the ΔΔC_t_ method as described earlier (Kim *et al.*, 2016). The following primers were used to check the quantitative transcriptional level of Prkab1, Vim, and Ahsa1; 5’-TCCGATGTGTCTGAGCTGTC-3’ and 5’-CCCGTGTCCTTGTTCAAGAT-3’ for Prkab1 (Bandow *et al.*, 2015), 5’-TCCACACGCACCTACAGTCT-3’ and 5’-CCGAGGACCGGGTCACATA-3’ for Vim (Ulmasov *et al.*, 2013), 5’-CAGAGGGGCACTTTGCCACCA-3’ and 5’-CACGGCCTTCCATGCACAGCT-3’ for Ahsa1.

### 2.9 Statistical analysis

Data were analyzed using Prism 4.0 (GraphPad Software, San Diego, CA). Non-paired Student’s *t*-test was used when two groups were being compared. Differences were considered statistically significant for p-values <0.05. Data are represented as the mean ± S.E.M. of at least two separate experiments performed at least in triplicate.

## 3. Results

### 3.1 Acute exposure to H_2_S induces motor behavioral deficits and seizures in C57 black mice.

Locomotor activities of mice were assessed on days 2, 4, and 6. Results indicated that the horizontal and vertical activities were decreased by more than 50% and were statistically different in mice exposed to acute H_2_S compared to the breathing air group (Fig. 2 A and B). Total distance traveled was also decreased by more than 50% and was also statistically different in H_2_S-exposed mice compared to controls (Fig. 2 C). More importantly, mice exposed to H_2_S exhibited severe seizure activity. On day 1, on average, seizure activity was observed starting at 15 min of H_2_S exposure. More than 50% of mice were shown to have seizure activity at 40 min of exposure to H_2_S. Mice exposed to H_2_S more than once exhibited seizure activity after only 5 min of H_2_S exposure. In these groups of mice, at 10 min of exposure to H_2_S, 40% of mice exhibited seizure activity on day 6. Collectively, these results indicate increased susceptibility of mice to H_2_S after each successive H_2_S exposures, suggesting that the toxic effects of H_2_S exposure are cumulative (Fig. 2D).

**Figure 2.**
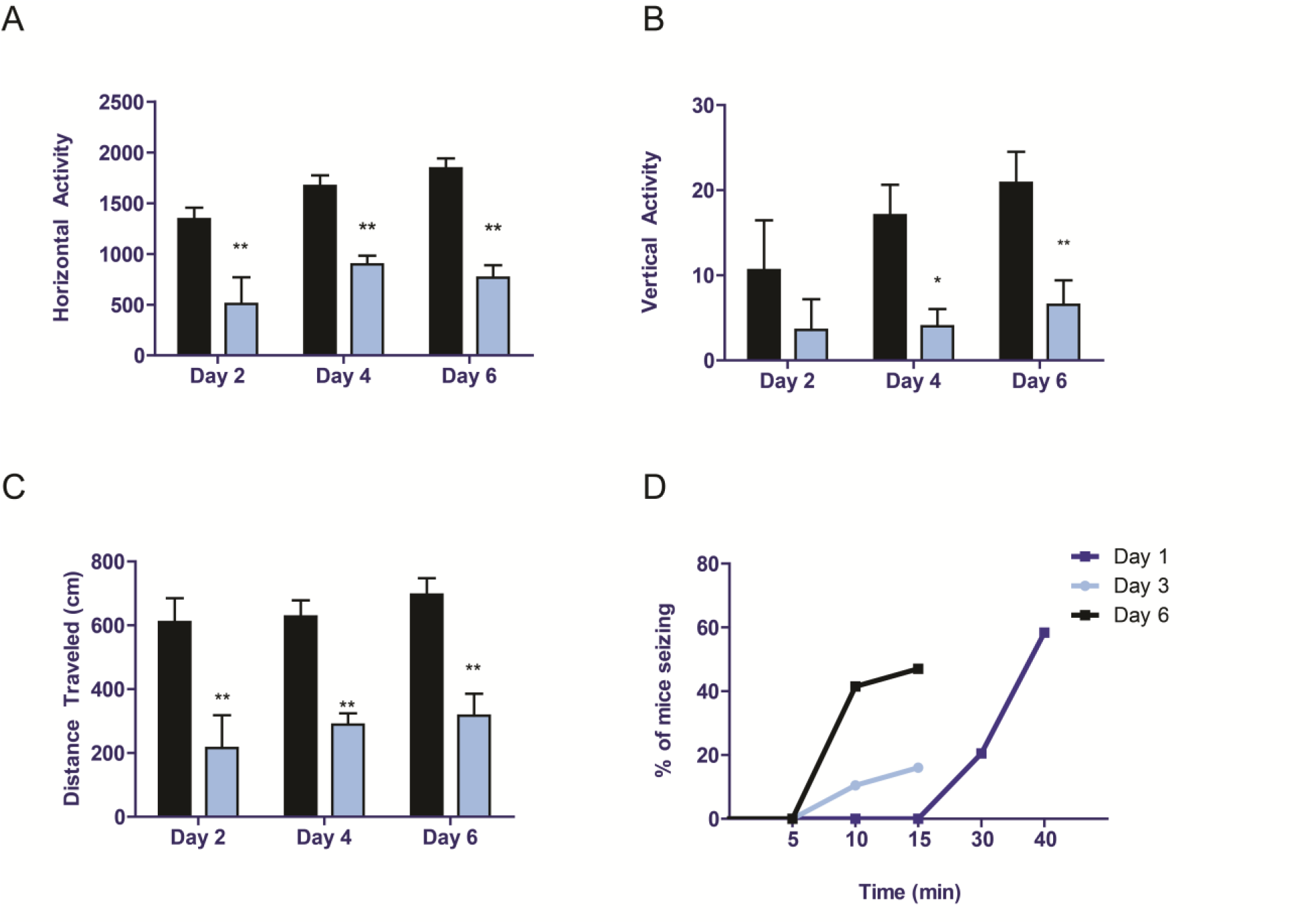
Acute exposure to hydrogen sulfide induced motor behavioral deficits. C57 black mice were exposed to H_2_S as shown in Fig. 1. Locomotor activity was measured using an automated VersaMax locomotor activity monitor during 10 min time test on days 2, 4, and 6. Horizontal activity (A), vertical activity (B), and total distance traveled (C) were analyzed between groups. Seizure activity was analyzed on a time scale (D). Asterisks (*, p < 0.05; **, p < 0.01) indicate statistically significant differences between H_2_S and breathing air negative control groups. (One and half column figure)

### 3.2 Acute hydrogen sulfide exposure induces brain damage

Motor behavioral deficits induced by exposure to H_2_S may be a sign of injury to the central nervous system (CNS). Therefore, histopathology was performed to identify the effects of H_2_S exposure in the IC. Mice were exposed to H_2_S as designated in Fig. 1 and sacrificed at multiple time points (day1, 3, or 7) for this portion of study. Microscopic examination of PFA-perfused brains revealed H_2_S-induced neurodegeneration and loss of neurons in the IC (Figure 3A). By day 7, mice exposed to H_2_S exhibited severe damage to the IC, with necrosis, vacuolar change, and infiltration by neuroglial cells. In order to further characterize loss of neurons, IC tissues were analyzed by immunohistochemical staining for the neuron specific marker, NeuN. On day 7, there was a marked loss of neurons in the IC of H_2_S-exposed mice, compared with the breathing air group. Enumeration of neurons in the IC revealed approximately 70 % loss of IC neurons by day 7 of H_2_S exposure (Fig. 3 B and C) with infiltration by glial cells. In H_2_S exposed mice, neurons in the midbrain adjacent to the IC appeared unaffected morphologically and glial cell numbers and activation state were not altered. These results demonstrate selective loss of neurons of the IC.

**Figure 3.**
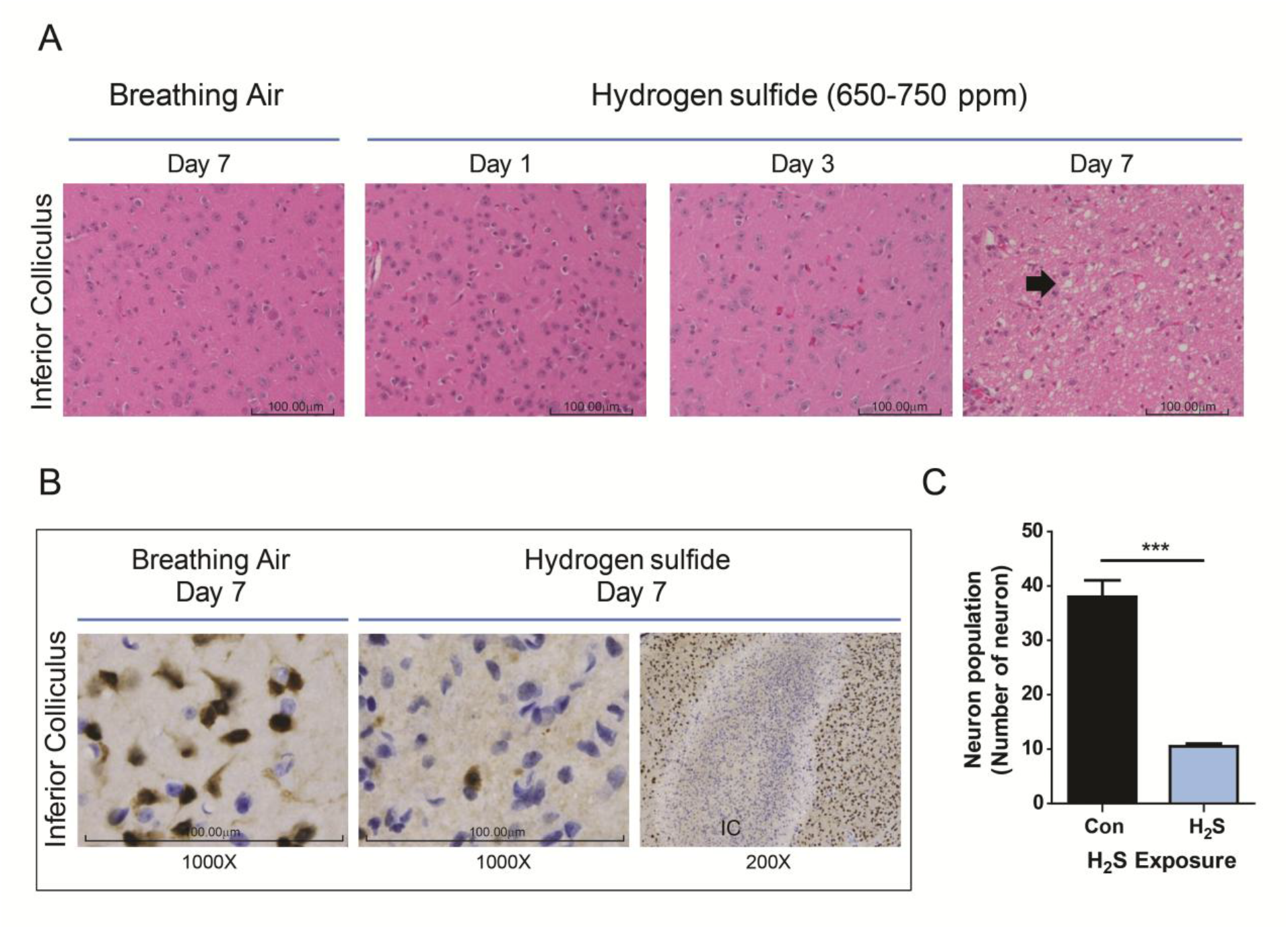
Neurodegeneration and necrosis in the IC of mice exposed to 650-750 ppm H_2_S in acute short-term repeated inhalation exposures over 7 days. Note the loss of neurons and development of clear vacuoles in the neuropil (arrow) on day 7 in mice exposed to H_2_S and the minimal morphologic effects to earlier H_2_S exposures or breathing air (A). Inhalation of H_2_S resulted in marked and selective loss of neurons in the IC (1000X magnification images, B), with retention of neurons in the regions surrounding the IC (200X magnification image). Computer aided image analysis of NeuN immunostained sections reveals marked loss of neurons in the IC of mice exposed to H_2_S (p=0.001, Students t test). Representative photomicrographs of mice exposed to breathing air or H_2_S, hematoxylin and eosin (A) and NeuN immunohistochemistry (B), 400X magnification (A) and 1000X and 200X magnification (B). Neurons in the IC region of the H_2_S exposed mice on day 7 were enumerated and compared to breathing air control (C). (Two column figure)

### 3.3 Changes in the Broad Spectrum Proteome following acute H_2_S Exposure

Proteomic profiles of the IC were determined using TMT peptide labeling coupled with LC/MS/MS analysis by comparison of the breathing air negative control group and the H_2_S exposed group is described in the methods section. Mass spec was able to identify 598, 562, and 546 altered proteins for days1, 2, and 4 of H_2_S exposure, respectively. Subsequent analyses showed alterations of protein expressions (Fig. 4, heatmap) with 36, 12 and 13 proteins being significantly downregulated (below 0.83 fold) and 8, 6, and 12 proteins being significantly upregulated (above 1.2 fold) compared to the control for days 1, 2 and 4 of H_2_S exposure, respectively. A single (day 1) H_2_S exposure, in which mice were euthanized only 2 h post a single exposure, demonstrated the highest range of fold protein expression changes compared to the control with the majority of proteins being in the downregulation cluster. This was followed by day 2 exposures with equal distribution of upregulated and downregulated proteins, while day 4 manifested the lowest range of fold changes with the least changes in the proteomic profile (Fig. 5 A, B, C, D, scatter plots).

**Figure 4.**
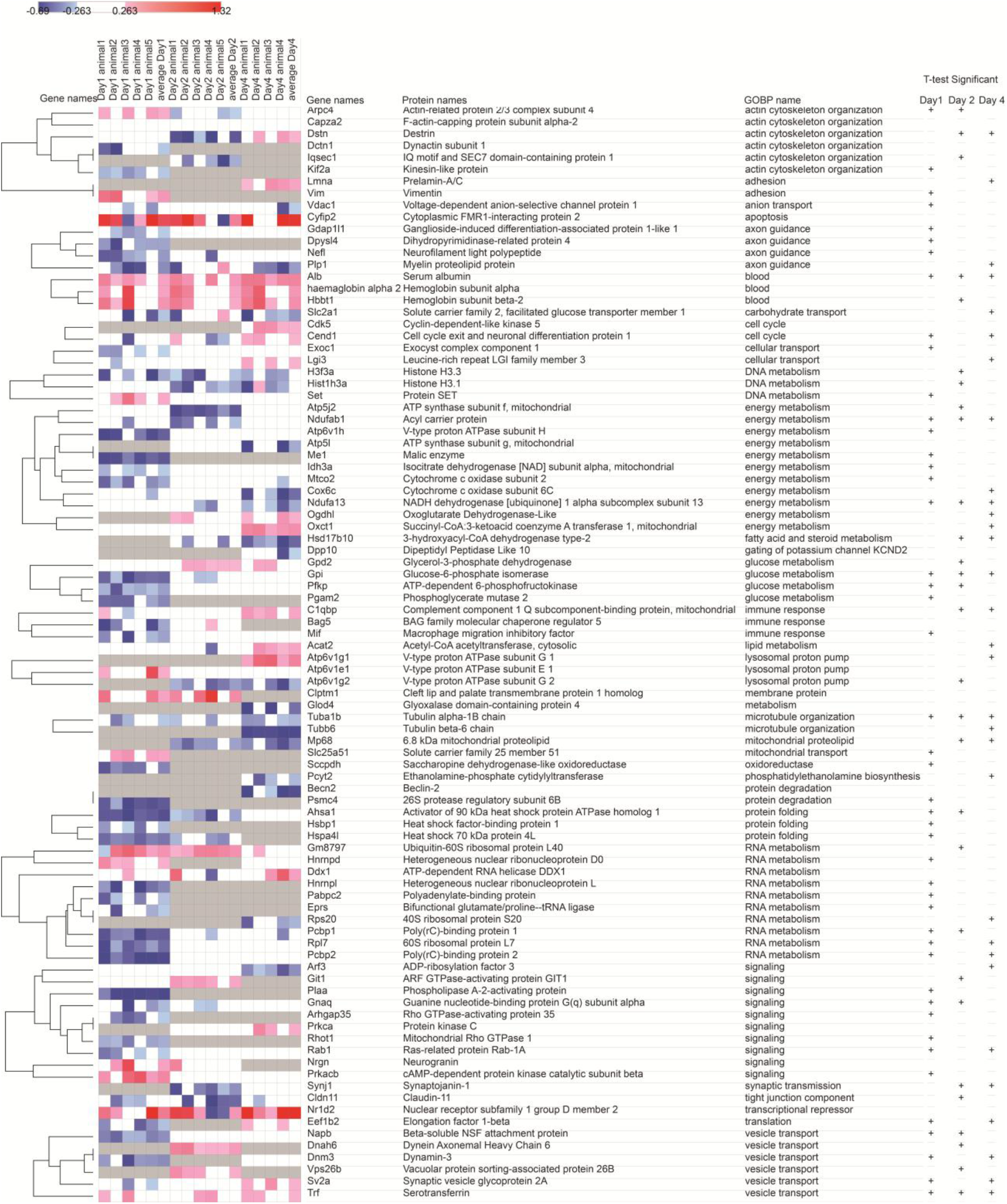
Heatmap of IC changes in proteomic profile following H_2_S exposure Morpheus rendered Heatmap of determined changes in protein expression on days 1, 2 and 4. H_2_S exposure using hierarchical clustering by Euclidean distance, row average linkage and grouping of rows by gene ontology for biological process (GOBP). Heatmap displays fold expression values of five (day 1 and day 2) or four (day 4) IC tissue samples, and average fold expression values, mouse gene identifiers (gene names), protein names, and GOBP. “+” indicates t-test (of fold expression values) significantly modulated proteins; grey not identified, white non regulated, red upregulated (above 1.2 fold expression vs. control), blue downregulated (below 0.83 fold expression vs. control) converted into a log2 data display. (Two column figure)

**Figure 5.**
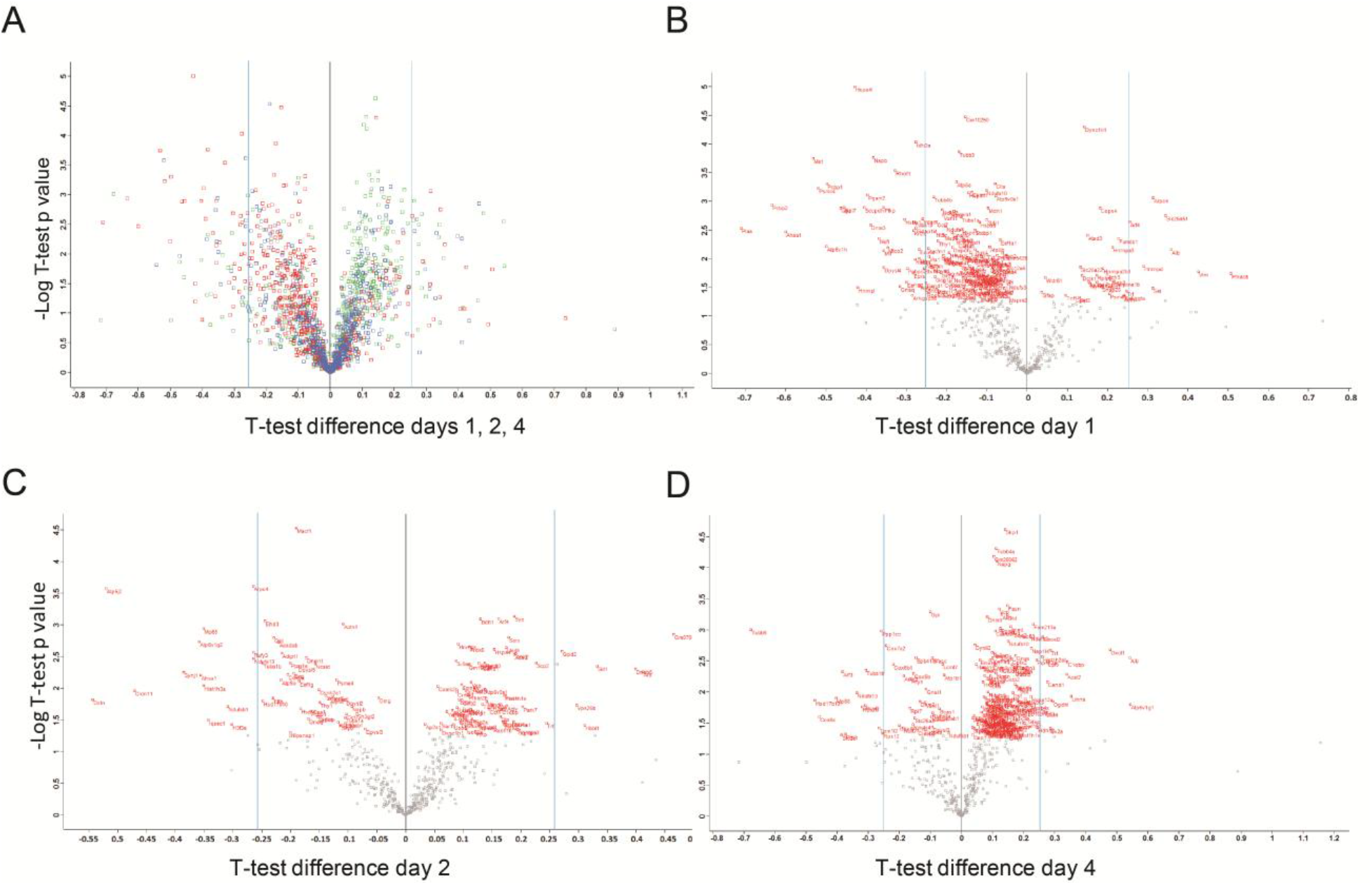
Overall distribution of IC changes in proteomic profile following H_2_S exposure A)Scatter Plot to display overall distribution of one-sample t-test differences in protein expression profiles vs. –Log one-sample t-test p values following day 1 (red), day 2 (blue), and day 4 (green) H_2_S exposure. Scatter Plots of one-sample t-test fold protein expression differences vs. –Log one-sample t-test p values from B) day 1, C) day 2, and D) day 4 of H_2_S exposure. One-sample t-test of significantly modulated proteins in the H_2_S exposure group vs. the control according to fold expression values are displayed with official mouse gene symbols in red. (Two column figure)

We further analyzed H_2_S-dependent alterations in the proteome using Perseus annotation for gene ontology (GO) biological pathways that were summarized into general categories when applicable by integrating gene cards information (www.genecards.org). We found that several gene ontology functions were affected after H_2_S exposure (Table 2). Acute exposure to H_2_S on day 1 resulted in alteration of RNA, energy and glucose metabolism, and changes in signaling pathways, axon guidance, actin cytoskeleton reorganization, and vesicle transport. On day 2 changes in gene ontology included glucose and energy metabolism, actin cytoskeleton organization and vesicle transport. Changes in energy and RNA metabolism were most abundant on day 4. Interestingly, there was an upregulation of cytoplasmic interacting protein Cyfip2 and nuclear hormone receptor and transcriptional repressor family Nr1d2 that were observed at all three time points. There was also a downregulation of anti-apoptotic and immunogenic protein Bcl2 associated athanogene Bag5 on day 1, suggesting potential induction of apoptosis. This is further supported by an observed downregulation of core histones H3f3a, and Hist1h3a that might result in a DNA damage response. Evidence of potential immunogenic changes was given by an upregulation in complement binding protein C1qbp on day 3 and a downregulation in antiinflammatory protein macrophage migration inhibition factor Mif on day 1. These latter changes suggest inflammation is involved in H_2_S-induced neurotoxicity.

**Table 2.** Functional annotation of significantly modulated protein expression in the IC following H_2_S exposure on days 1, 2, and 4. Table displays total number of proteins associated with GO-term definition. A gradient scheme was applied to better indicate number distribution with lower numbers in light orange and higher numbers in dark orange. Note that day 1 represents mice exposed once and euthanized 2 h post-exposure. (One and half column table)

| Gene Ontology Term | Day1 | Day2 | Day4 |
| --- | --- | --- | --- |
| actin cytoskeleton organization | 3 | 4 | 2 |
| adhesion | 1 | 0 | 1 |
| anion transport | 1 | 0 | 0 |
| axon guidance | 3 | 0 | 1 |
| blood | 1 | 2 | 1 |
| cell cycle | 1 | 0 | 1 |
| cellular transport | 1 | 0 | 1 |
| DNA metabolism | 1 | 2 | 0 |
| energy metabolism | 6 | 3 | 5 |
| glucose metabolism | 3 | 3 | 1 |
| immune response | 1 | 1 | 1 |
| microtubule organization | 1 | 1 | 2 |
| mitochondrial transport | 1 | 0 | 0 |
| oxidoreductase | 1 | 0 | 0 |
| protein degradation | 1 | 0 | 0 |
| protein folding | 3 | 1 | 0 |
| RNA metabolism | 7 | 2 | 3 |
| signaling | 6 | 2 | 2 |
| vesicle transport | 3 | 3 | 2 |
| fatty acid steroid metabolism | 0 | 1 | 1 |
| mitochondrial proteolipid | 0 | 1 | 1 |
| synaptic transmission | 0 | 1 | 1 |
| tight junction component | 0 | 1 | 0 |
| carbohydrate transport | 0 | 0 | 1 |
| lipid metabolism | 0 | 0 | 1 |
| phosphatidylethanolamine biosynthesis | 0 | 0 | 1 |
| translation | 0 | 0 | 1 |
| lysosomal proton pump | 0 | 1 | 1 |

### 3.4 Validation of proteomic changes of genes after H_2_S exposure

To confirm the observed changes in the IC proteomic profile following acute exposure to H_2_S, several genes were analyzed by quantitative RT-PCR. Mice were sacrificed 2 h after each designated time point to measure cellular response right after H_2_S exposure. Protein kinase AMP-activated non-catalytic subunit beta 1 (Prkab1, also called Ampk) is known to correlate with calcium fluctuations as a measure of cellular calcium response (Yong *et al.*, 2010). Prkab1 mRNA expression was analyzed as a means to measure the calcium-dependent cellular response following H_2_S exposure. Prkab1mRNA expression was upregulated following H_2_S exposure on day 1, 2 and 4 which is in line with the mass spec observed upregulation of protein Kinase cAMP-activated catalytic subunit alpha (Prkca) and calcium activated protein kinase C beta (Prkacb) on exposure day 1. To confirm potential H_2_S exposure dependent pro-inflammatory and ischemic effects, mRNA expression of Vimentin (Vim) was measured. Vim mRNA expression demonstrated a steady increase throughout all days of H_2_S exposure (Fig. 6 B). This is further supported by the increased protein expression of Vim on exposure day 1, while mass spec lacked detection of Vim in the other samples.

**Figure 6.**
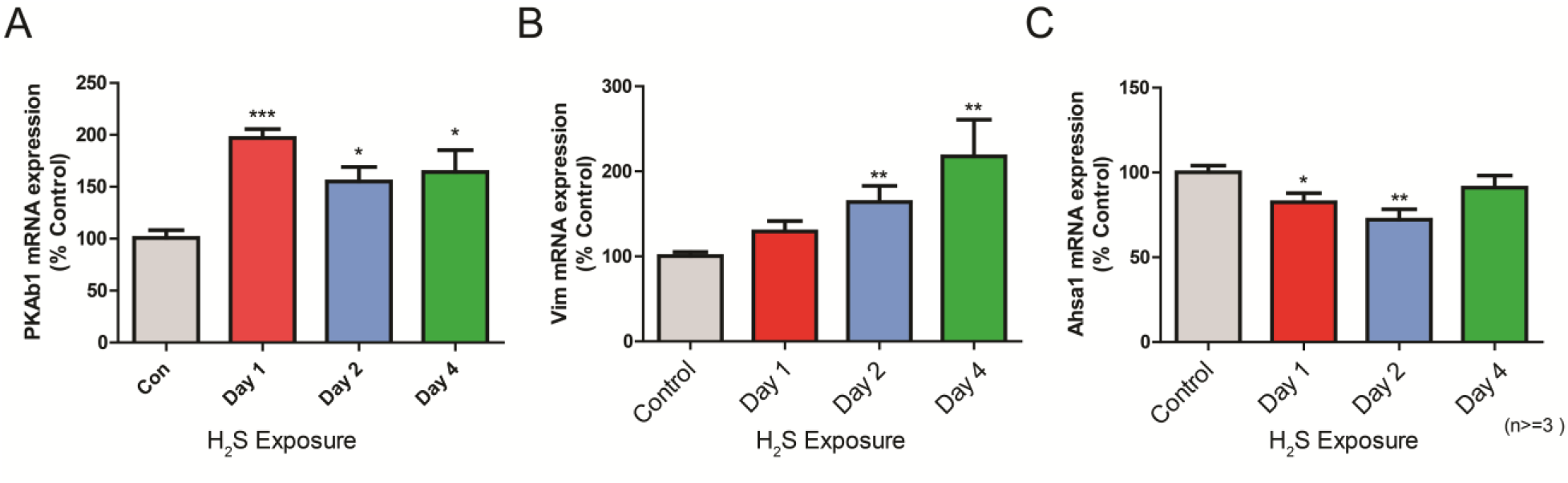
Validation of expression changes of three genes at the mRNA level in IC after H_2_S exposure. The transcriptional level of three genes (A; Prkab1, B; Vim, C; Ahsa1) was measured by quantitative PCR. Samples were normalized with the reference gene Gapdh. Data are represented as the mean ± S.E.M. Note that only a single acute exposure to H_2_S was needed to cause upregulation of gene expression of PkAb1 and Vim or downregulation of Ahsa1 gene expression. (Two column figure)

Due to a mass-spec observed downregulation in the activator of 90 kDa heat shock protein and chaperone ATPase homlog1 (Ahsa1) on day 1 and 2, mRNA expression of Ahsa1 was examined by RT-PCR (Fig. 6 C). Similar to the mass spec observed protein expression, mRNA expression of Ahsa1 demonstrated a significant 30-40 % downregulation on days 1 and 2 of H_2_S exposure, compared to the breathing air control group.

## 4. Discussion

Acute H_2_S exposure often leads to neurological sequelae among human survivors, suggesting a trigger of neurodegeneration cascade following acute H_2_S exposure (Matsuo *et al.*, 1979; Tvedt *et al.*, 1991a; Tvedt *et al.*, 1991b; Kilburn, 1993; Snyder *et al.*, 1995; Woodall *et al.*, 2005). However, the exact cellular and molecular mechanisms underlying delayed neurological disorders after acute H_2_S exposure are poorly understood. Few studies have been done to characterize mechanisms of H_2_S exposure-induced neurotoxicity in survivors of acute H_2_S poisoning. We recently developed an inhalational mouse model which recapitulates H_2_S-induced neurotoxicity observed in humans (Anantharam *et al.*, 2017). In this model, mice exposed to H_2_S showed clinical and pathological characteristics as shown in human patients. In case studies of human H_2_S inhalation exposure, H_2_S concentrations > 500 ppm were reported to rapidly induce severe neurotoxicity and temporary paralysis of breathing, leading to immediate collapse (Guidotti, 1996; Chou *et al.*, 2016). Survivors of acute high H_2_S exposures (> 500 ppm) developed chronic neurological problems such as loss of memory, loss of control of facial muscles, persistent headaches, and neurodegeneration. These patients also exhibited motor behavioral deficits such as spastic gait (Wasch *et al.*, 1989; Tvedt *et al.*, 1991a; Tvedt *et al.*, 1991b; Callender *et al.*, 1993). Brain scan images of patients accidently exposed to H_2_S suggested a dysregulation of basal ganglia region that governs motor behavior (Schneider *et al.*, 1998; Nam *et al.*, 2004). This is consitent with the observed motor deficits in our RotaRod studies that are in agreement with clinical phenotypes of H_2_S exposed human patients. According to Wasch *et al.*, the brainstem was demonstrated to accumulate the highest concentration of H_2_S compared to other brain regions in the rat (Wasch *et al.*, 1989). We have previously identified and herein confirmed that the IC, a brainstem region, is highly sensitive to H_2_S-induced necrosis.

The relative sensitivity of the IC to H_2_S-induced injury reflects the unique elements and cell populations in this region of the brain. We observed histologic features of neurodegeneration, neuronal death, and reactive gliosis is in the IC starting on day 3 with massive cell necrosis and glial scarring in the IC on day 7. Immunohistochemistry with neuron specific antibody (NeuN) revealed severe neuronal loss in the IC with H_2_S exposure. Loss of neuronal cells was more than 70 % compared to breathing air control group in IC. In contrast, brain stem regions adjacent to the IC were shown to be relatively unaffected, indicating unique susceptibility of the IC region to H_2_S toxicity. The function of the IC is to integrate auditory and other sensory signals and is reportedly a region with high metabolic rate requirements (Ridgway *et al.*, 2006; Houser *et al.*, 2010) while it is also the most highly vascularized brain region (Gonzalez-Lima *et al.*, 1997). H_2_S is a systemic metabolic toxicant that reportedly interferes with ATP synthesis. It is reasonable to infer that brain regions with high blood supply and metabolic rates such as the IC are more vulnerable to H_2_S-induced toxicity.

One of the hypotheses underlying H_2_S-induced neurotoxicity is that H_2_S causes hypoxia-dependent neurodegeneration following H_2_S induced hypotension and pulmonary edema-dependent oxygen deprivation in inhalation victims of H_2_S poisoning (Nicholls and Kim, 1982; Dorman *et al.*, 2002; Miyazato *et al.*, 2013; Rumbeiha *et al.*, 2016). However, the proximate molecular pathways by which these events lead to neurodegeneration are unknown. In this study, we used proteomic analysis to define cellular and molecular mechanisms of H_2_S induced neurodegeneration during early stages of injury. We demonstrated that H_2_S exposure induced significant alteration of protein expression in the IC in various biological pathways, including cellular morphology, energy metabolism, and calcium signaling. It was most interesting that a single H_2_S exposure exerted the most significant changes, suggesting a single exposure, as commonly occurs during single accidental H_2_S exposures, is sufficient to trigger such proteomic changes.

Energy failure is another hypothesis of H_2_S-induced neurotoxicity because it exerts its cellular toxicity by inhibition of cytochrome C oxidase which plays a crucial role in mitochondrial ATP synthesis (Nicholls and Kim, 1982; Dorman *et al.*, 2002). In this study, expression levels of cytochrome C oxidase-related proteins were not changed, while cytochrome C oxidase subunit 6 C (Cox6c) and mitochondrial cytochrome C oxidase subunit 2 (mt-Co2) were decreased on day 4 and 1, respectively, supporting potential impairment of energy production following H_2_S exposure. Several other proteins related to energy production were also altered. For instance, ATP synthase subunits Atp5j2 and ATP5l together with cytochrome C oxidases 6C were downregulated on days 2, and 4, respectively. In addition, citric acid cycle proteins malic enzyme (Me1) and phosphoglycerate mutase (Pgam2) were downregulated on day 1. Further studies are warranted to check whether Coxy6c and mt-Co2 are the bona fide targets of H_2_S toxicity. Collectively, these results suggest hypoxia, resulting in low oxygen tissue delivery, and impaired ATP synthesis arising from dysregulation of cytochrome C related proteins, likely work in concert leading to neuronal death in the IC.

The IC undergoes severe degenerative changes during H_2_S exposure including widespread neuronal loss, which is regionally selective. Long-term exposure to H_2_S was demonstrated to result in neuronal structural damage by demyelination (Solnyshkova, 2003). Interestingly, claudin-11 (Cldn11), a major tight junction component in neuronal myelin structures of the central nervous system (CNS) was downregulated following day 1 and 2 of H_2_S exposure supporting potential H_2_S induced neurodegeneration (Gow *et al.*, 1999; Tiwari-Woodruff *et al.*, 2001). Besides, mass spectrometry detected alterations in proteins related to cellular morphological changes included regulators of actin cytoskeleton and microtubule organization. Dynamine-3 (Dnm3), a GTP binding protein associated with the microtubule system was downregulated on day 1. Dynein Axonemal Heavy Chain 6 (Dnah6), a component of microtubule-associated motor system, was upregulated on day 4. Moreover, a couple of adhesion proteins were altered, e.g., vimentin (Vim) was upregulated on day 1 while prelaminin-A/C (Lmna) was upregulated on day 4 of H_2_S exposure. Dihydropyrimidinase-related protein Dpysl4, and neurofilament light polypeptide Nefl were downregulated on day 1 together with a downregulation of myelin proteolipid protein Plp1 throughout the study period. With regards to other cytoskeletal proteins, Tuba1b and Tubb6 were downregulated on day 4. These changes may impact cell integrity and function in the IC, leading to neuronal death.

H_2_S exposure has previously been shown to induce dysregulation of calcium concentrations (Garcia-Bereguiain *et al.*, 2008). It was further reported by others that exposure to H_2_S may affect regulation of calcium (Nagai *et al.*, 2004; Garcia-Bereguiain *et al.*, 2008; Yong *et al.*, 2010). In this study, the calcium binding protein neurogranin (Nrgn) demonstrated a nonsignificant upregulation on day 1. However, we found a significant upregulation in cAMP-dependent protein kinase that matched the expression pattern of Prka1 mRNA supporting potential activation of calcium-dependent signaling. These results agree with previous reports that demonstrated H_2_S-dependent calcium and cyclic-AMP signaling through Prka (Kimura, 2000; Yong *et al.*, 2010). Collectively, these results suggest calcium dysregulation as a potential mechanism of H_2_S-induced neurotoxicity.

Changes in immunogenic biomarkers represent one of earliest responses to cytotoxicity. Evidence of potential H_2_S-dependent immunogenic changes was given by a downregulation in anti-inflammatory protein macrophage migration inhibition factor Mif on day 1 and an upregulation in the reactive oxygen species (ROS) response and complement binding protein C1qbp on day 4. Immunogenic changes in MIF have been previously observed (Roger *et al.*, 2003). ROS response and complement binding protein C1qbp has been previously reported (McGee and Baines, 2011). In addition, there was an upregulation of cytoplasmic interacting protein (Cyfip2) and nuclear hormone receptor and transcriptional repressor family (Nr1d2) that were observed throughout the exposure period. Cyfip2 was previously shown by others to have a p53-response element in its promoter region and to be one of the direct targets of p53 (Jackson *et al.*, 2007). Besides, the anti-apoptotic and immunogenic protein Bcl2 associated athanogene (Bag5) was downregulated on day 1. Endoplasmic reticulum stress-induced downregulation of Bag5 has previously reported (Bruchmann *et al.*, 2013; Gupta *et al.*, 2016). Further evidence of potential cell stress-dependent impairment in protein folding was shown by an RT-PCR and mass spec detected downregulation in Ahsa1 that plays a role as chaperone and activating heat shock protein 90 (Okayama *et al.*, 2014; Tripathi *et al.*, 2014). These results indicate that inflammation may play an important role in H2S-induced neurotoxicity.

H2S-induced neurotoxicity has been suggested to resemble the injury caused by ischemic hypoxic conditions (Doujaiji and Al-Tawfiq, 2010; Rumbeiha *et al.*, 2016). Previous studies have suggested neurofilament Nefl, and collapsin Response Mediator Protein 3 (Dpysl4) as biomarkers of hypoxia and cerebral ischemia (Hou *et al.*, 2006; Lian *et al.*, 2015). Dpysl4 is cleaved by activated calpain reaction in response to cerebral ischemia (Hou *et al.*, 2006). Both Nefl and Dpysl4 were downregulated during H_2_S exposure. In addition, mass spec detected downregulation of phosphoglycerate mutase 2 (Pgam2), an ischemia biomarker, on day 1. This biomarker is suggested to correlate with the ischemic conditions (Li *et al.*, 2012). Vimentin (Vim) has been shown as another biomarker for ischemia (Li *et al.*, 2008). Mass spec also detected a steady increase in protein and mRNA expression of the inflammatory and ischemia biomarker Vim for the entire duration of the study. These data support ongoing progression of H_2_S-dependent ischemia-like conditions during acute H_2_S exposures, further supporting the role of hypoxia as a mechanism of H_2_S-induced neurotoxicity.

## 5. Conclusion

In this study we examined the early effects of acute H_2_S exposure on brains of mice following either a single H_2_S exposure or up to 4 short-term acute exposures. Taken together, acute exposure to H_2_S induced neurotoxicity, which manifested as progressive behavioral deficits. The IC was the most sensitive brain region, confirming our previous observations. Results point to demonstrated modulation of several proteome-based biological pathways in the IC. Specifically, results of proteomic and gene expression studies suggest calcium dysregulation, immune mediated inflammatory response, pro-apoptosis mechanisms, and ischemia-like cytotoxicity to be involved in H_2_S-induced neurotoxicity. This proteomic data provided important clues on mechanisms of H_2_S-induced neurological toxicity. Further research is recommended to better understand the singular or collective role of these potential mechanisms in H_2_S-induced mortality, neurotoxicity, and evolution of neurological sequelae.

## Acknowledgment

The authors do not have any conflict of interest. This work was partially supported by the Iowa State University College of Veterinary Medicine Seed grant, Startup funds and incentive account funds for Rumbeiha.

